# Exercise enhances hippocampal-cortical ripple interactions in the human brain

**DOI:** 10.1101/2023.05.19.541461

**Authors:** Araceli R. Cardenas, Juan F. Ramirez-Villegas, Christopher K. Kovach, Phillip E. Gander, Rachel C. Cole, Andrew J. Grossbach, Hiroto Kawasaki, Jeremy D.W. Greenlee, Matthew A. Howard, Kirill V. Nourski, Matthew I. Banks, Michelle W. Voss

## Abstract

Physical exercise acutely improves hippocampus-dependent memory. Whereas animal studies have offered cellular- and synaptic-level accounts of these effects, human neuroimaging studies show that exercise improves hippocampal-cortical connectivity at the macroscale level. However, the neurophysiological basis of exercise-induced effects on hippocampal-cortical circuits remains unknown. Experimental evidence supports that hippocampal sharp wave-ripples (SWR) play a critical role in learning and memory. Coupling between SWRs in the hippocampus and neocortex may reflect modulations in inter-regional connectivity required by mnemonic processes. Here, we examine the hypothesis that exercise modulates hippocampal-cortical ripple dynamics in the human brain. We performed intracranial recordings in epilepsy patients undergoing pre-surgical evaluation, during awake resting state, before and after an exercise session. Exercise increased ripple rate in the hippocampus. Exercise also enhanced the coupling and phase-synchrony between cortical ripples in the limbic and the default mode (DM) cortical networks and hippocampal SWRs. Further, higher heart rate during exercise, reflecting exercise intensity, was related to a subsequent increase in resting state ripples across specific cortical networks including the DMN. These results offer the first direct evidence that a single exercise session elicits changes in ripple events, a well-established neurophysiological marker of mnemonic processing. The characterization and anatomical distribution of the described modulation points to hippocampal ripples as a potential mechanism by which exercise elicits its reported short-term effects in cognition.

## Introduction

Physical exercise improves memory and learning in rodents and humans^1,2^. Recent data highlight the importance of hippocampal circuits to these cognitive benefits^3,4^. Animal studies have focused on the molecular, histological and endocrine mechanisms by which exercise drives synaptic plasticity and neurogenesis in the hippocampus^5–7^. Human neuroimaging studies have shown that exercise increases hippocampal-cortical and medial temporal lobe functional connectivity in the Default Mode Network (DMN) both acutely^8,9^ and after months of training^4,10,11^. However, all human data to date rely exclusively on hemodynamic measures of hippocampal function, which in the scale of seconds, offers only indirect inference about neural function. For instance, at a sluggish pace, the blood oxygen level-dependent (BOLD) signal cannot speak to the millisecond circuit-level changes underlying functional connectivity and ultimately cognition. Moreover, differences in hippocampal and neocortical neural dynamics between rodents, non-human and human primates have been a matter of debate. In regard to exercise, rodent studies are limited in the translation of exercise intensity based on their unique vascular physiology^12^. Examining the acute effects of exercise on electrophysiological measures of neural function, with direct recordings from the human brain, offers a rare opportunity to bridge animal and human findings, resolve millisecond-scale circuit dynamics, and advance understanding of how exercise influences human brain function.

There is a growing understanding of the neurophysiological mechanisms supporting hippocampal-dependent learning and memory^13^. For instance, during periods of quiescence or non-rapid-eye-movement (NREM) sleep, slow field potential deflections accompanied by high frequency oscillations (HFOs), known as sharp wave-ripple (SWR) complexes, occur in the hippocampus and play a critical role in memory consolidation^14,15^.Occurrence of hippocampal SWR is linked to widespread patterns of cortical activation and simultaneous subcortical deactivation^16,17^. Furthermore, the occurrence of ripples in the neocortex^18^ and their functional role^19–21^ has gained attention recently. In both rodents and humans, the coupling between hippocampal and neocortical ripples has been functionally associated with memory retrieval and learning^19–21^. Overall, recent evidence indicates that hippocampal-cortical ripple coupling is a neurophysiological marker of transient functional connectivity^22^.

These findings invite the hypothesis that exercise modulates ripple dynamics in the hippocampus, medial temporal lobe, and cortical networks such as the DMN. Here, we obtained intracranial electroencephalographic (iEEG) recordings in patients with drug resistant epilepsy undergoing pre-surgical evaluation using stereo and subdural EEG, during awake resting state sessions, before and acutely after physical exercise. Our central hypothesis was that exercise would acutely increase ripple rate in the hippocampus, and enhance hippocampal-cortical interactions in the ripple frequency band (70-160 Hz).

Our results confirm and extend those predictions. Across patients, exercise selectively enhanced ripple-associated hippocampal connectivity with the limbic (LIM) network and the DMN, as evidenced by enhanced ripple coupling and ripple phase synchrony. Crucially, higher heart rates during exercise, reflecting higher exercise intensity, predicted greater enhancement of resting state ripple dynamics in specific neural networks such as the DMN.

Overall, we show that a single session of light to moderate intensity physical exercise triggers changes in human ripple hippocampal-cortical dynamics. The characterization and anatomical distribution of those changes, predominant in limbic and DM networks, advances the possibility that these modulations account for the previously described effects of exercise on learning and memory.

## Materials and methods

### Participants

A total of 17 patients with drug resistant epilepsy undergoing pre-surgical evaluation using stereo and subdural EEG, completed the acute exercise paradigm, all while being monitored with chronic intracranial EEG recordings. Two of these patients were excluded from the analysis because of severe anatomical anomalies (antecedents of intrauterine stroke and previous left ATL). Another patient was excluded because of reported and evident severe cognitive deficits. Patient demographics and seizure onset zone for the remaining 14 patients are described in Table 1. Research protocols aligned with best practices recently aggregated in Feinsinger et al.^23^ The University of Iowa Human Subjects Review Board approved the research protocol and written informed consent was obtained from each subject before the study.

**Table 1.** Patient Demographics. Included patients experimental IDs (N=14), accompanied by their age (mean, median, and range are provided at the bottom of the table); sex (M: Male; F: Female; 4 females, 10 males); anatomical location of the seizure onset zone; and average heart rate as a percentage of the age-related maximum (mean, median, and range are provided at the bottom of the table). RPE: Ratings of Perceived Exertion. *NA*: RPE for one participant was not available due to missing self-report during exercise. PHG: Parahippocampal gyrus; ITG: Inferior temporal gyrus.

| Patient | Age | Sex | Seizure focus | Avg HR<br>(% max) | Avg RPE<br>(6-20) |
| --- | --- | --- | --- | --- | --- |
| S01 | 30 | M | L ITG including PHG | 62 | 11 |
| S02 | 18 | M | R parietal med | 59 | 12 |
| S03 | 50 | F | L ant mesial temporal | 59 | 12 |
| S04 | 18 | M | R frontal ant | 45 | 13 |
| S05 | 29 | M | L insula lesion | 52 | 12 |
| S06 | 28 | F | R frontal ant | 54 | 15 |
| S07 | 20 | M | R orbitofrontal | 56 | 11 |
| S08 | 33 | M | L mesial temporal | 62 | 11 |
| S09 | 17 | M | R sup and mid temporal | 60 | 11 |
| S10 | 22 | F | R superior temporal | 51 | 10 |
| S11 | 19 | M | L frontal | 68 | 11 |
| S12 | 32 | M | L temporal (fusiform,<br>lingual) | 59 | 15 |
| S13 | 25 | F | R temporoparietal | 59 | *NA* |
| S14 | 18 | M | L mesial temporal | 62 | 11 |
| **N** | **14** | **10 M** |  |  |  |
| **Mean** | **25.6** |  |  | **57.7** | **11.9** |
| **Median** | **23.5** |  |  | **58.9** | **11** |
| **Range** | **17-50** |  |  | **45-68** | **10-15** |

### Acute exercise paradigm

Prior to the acute exercise paradigm, all participants were screened using the Physical Activity Readiness Questionnaire (PAR-Q, Canadian Society for Exercise Physiology). Weekly physical activity was also assessed using the Paffenbarger Physical Activity Questionnaire (PPAQ). After a first rest period of approximately 20 minutes, participants had a guided exercise session on a MagneTrainer-ER mini-bike (MagneTrainer) placed on the ground beside their bed or at the foot of a chair next to the bed. This session was followed again by a post-exercise rest period of similar duration (Fig 1A).

**Figure 1.**
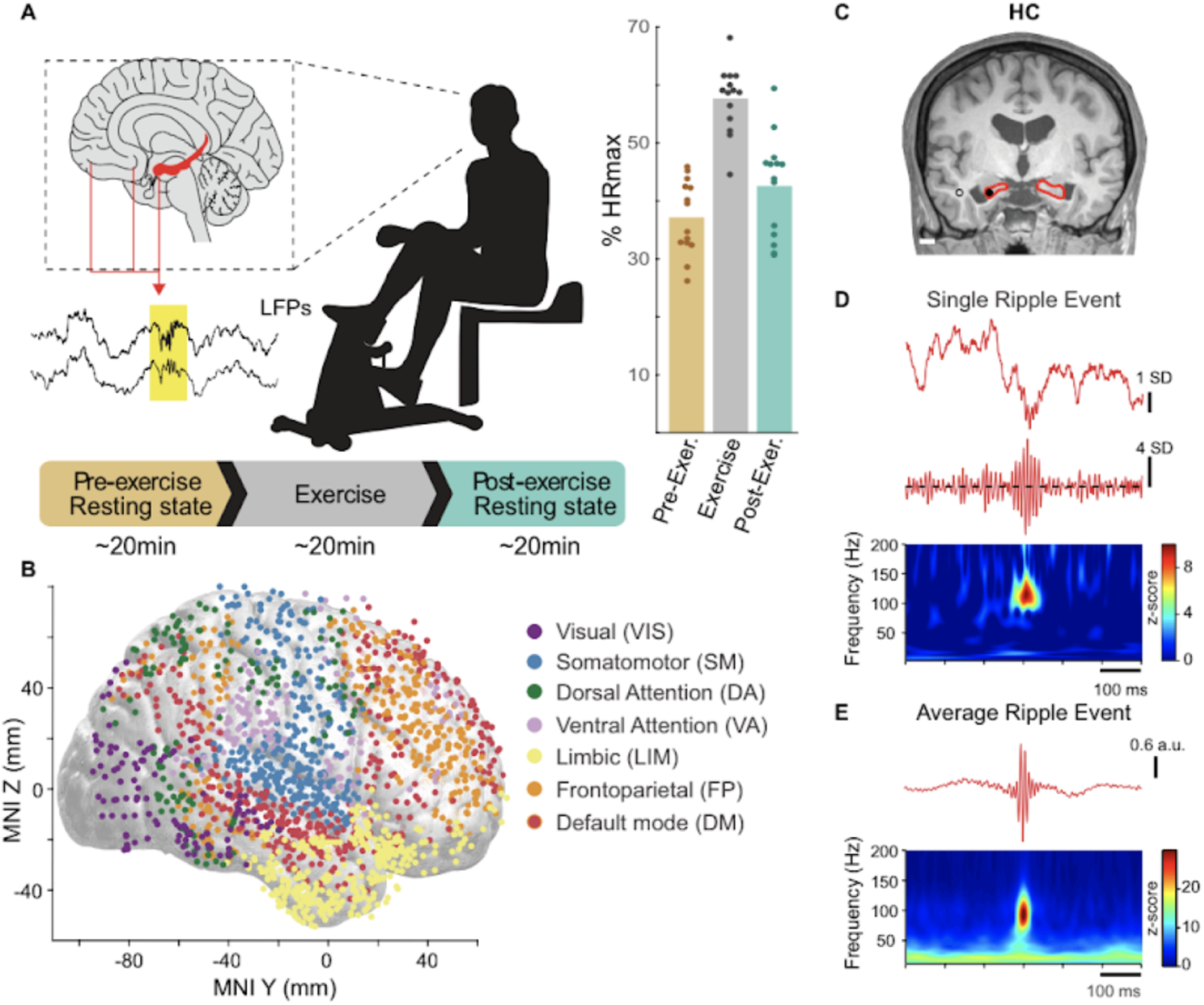
Paradigm, electrode coverage and ripple detection. (**A**) Left: Illustration of the experimental paradigm. Neurosurgical patients (a total of 14 included) were instructed to complete an acute exercise paradigm while intracranial EEG activity was continuously recorded. A pre-exercise resting period (∼20 min), was followed by exercise on a mini-bike (∼20 min), then followed by a postexercise resting period (∼20 min). Right: pre-, during, and post-exercise heart rate (percentage of age-predicted maximum) showing that compliance to the acute exercise protocol was satisfactory (50-60% target range) across subjects (individual dots). \**p*<0.001 paired-samples t-test for the comparison between pre-exercise resting state HR% and exercise HR%. (**B**) Cortical electrode coverage across participants. Contacts were grouped into 7 canonical networks using the Yeo-parcellation according to their anatomical location, as indicated by colors (visual, somatomotor, dorsal attention, ventral attention, limbic, frontoparietal, and default-mode). **(C)** One exemplary contact (filled circle) located in hippocampus shown overlayed in anatomical MRI T1 image. The nearest white matter contact (empty circle in the MRI image) was used for re-referencing. **(D)** Raw (<300 Hz) and filtered (70-160 Hz) iEEG (top/middle), together with the spectral decomposition of the raw iEEG (bottom), show that ripple events can be reliably identified in the electrical activity of the hippocampus. **(E)** Top: Single-channel first principal component of the peri-ripple iEEG traces (<300 Hz); Bottom: Averaged z-scored peri-ripple spectrogram.

Participants were explained the structure of the whole paradigm before starting the recordings. They were specifically instructed to lie on the bed with eyes closed after finishing the exercise session. Participants were asked to relax and avoid talking during the pre and post exercise resting state periods. An experimenter in the room documented if the experimental conditions were not satisfied. Audio recordings were used to further verify the start and end of the resting state periods to be analyzed.

In the exercise session, participants started with a 5-minute warm-up and then cycled at a light-to-moderate intensity of 50-60% of their age-predicted maximum heart rate (HR). Participants were given their target HR zone, so that they could monitor their pacing and adjust effort as needed. At the start of the warm-up period, bike resistance was set to half of the maximum resistance, and all participants were asked to start pedaling at a rate of 15 mph. Resistance was then adjusted until reaching a level that increased HR to the target HR zone and which the participants thought they could maintain for 20 minutes. Ratings of perceived exertion (RPE 6-20) were recorded following the warm-up and at 10-minute increments during exercise. The ECG read-out for HR was documented by the experimenter at the start of the pre- and post-exercise rest periods. Average RPE and relative HR (%HRmax) for each participant during exercise is shown in Table 1.

### Recording of iEEG data

Recordings were made using subdural and depth electrodes, manufactured by PMT Corporation, Chanhassen, MN (participants S01-S09) or Ad-Tech Medical, Racine, WI (all other participants). Electrodes were implanted solely on the basis of clinical requirements, as determined by a team of epileptologists and neurosurgeons. Details of electrode implantation, recording and iEEG data analysis have been previously described in detail ^24–26^. In brief, subdural arrays consisted of platinum-iridium discs (2.3 mm diameter, 5–10 mm inter-electrode distance), embedded in a silicon membrane. These arrays provided coverage of frontal, parietal, temporal, and occipital cortex over the convexity of the cerebral hemispheres. Depth arrays (8–12 electrodes, 5 mm inter-electrode distance) targeted the amygdala, hippocampus, cingulate, insular cortex and the superior temporal plane. A subgaleal electrode, placed over the cranial vertex near midline, was used as a reference in all participants.

In participants S01 through S09 data acquisition was controlled by a TDT RZ2 real-time processor (Tucker-Davis Technologies, Alachua, FL); in S10 through S14 data acquisition was performed using a Neuralynx Atlas System (Neuralynx, Bozeman, MT). Recorded data were amplified, filtered (0.1–500 Hz bandpass, 5 dB/octave rolloff for TDT-recorded data; 0.7–800 Hz bandpass, 12 dB/octave rolloff for Neuralynx-recorded data), digitized at a sampling rate of 2034.5 Hz (TDT) or 2000 Hz (Neuralynx) and stored for subsequent offline analysis.

### Anatomical reconstruction and ROI parcellation

All patients underwent whole-brain high-resolution T1-weighted structural magnetic resonance imaging (MRI) scans (1mm^3^ isotropic resolution) before electrode implantation. After electrode implantation, patients underwent MRI and thin-slice volumetric computerized tomography (CT) scans (1mm^3^ isotropic resolution). Locations of the depth and subdural electrode contacts were extracted from post-implantation MRI and CT scans, based on localized magnetic susceptibility artifacts and metallic hyperintensities, respectively. Anatomical locations estimated from postoperative scans were then projected onto preoperative MRI scans using non-linear three-dimensional thin-plate spline warping to account for geometric distortions, guided by manually selected control points throughout each individual brain (typically numbering 50-100) until satisfactory alignment was achieved, with a target residual displacement no greater than 2-3 mm. Data across participants were pooled by transforming the electrode locations into a standard coordinate space referenced to the Montreal Neurological Institute (MNI) 152 template. This was done for each participant using automated volumetric nonlinear co-registration to the MNI152 aligned CIT168 template brain ^27^. Volumetric co-registration used the ANTs software tool^28^.

Recording sites were assigned an anatomical label based on the Destrieux atlas implemented as individualized and automated parcellation of cortical gyri using the Freesurfer software package^29,30^. Further, cortical recording sites were also assigned a brain network label based on the Yeo7 network parcellation available in standard MNI space^31^ (Fig 1B). For depth electrodes, anatomical assignment was informed by MRI sections along sagittal, coronal and axial planes, and review from the Neurosurgical team (Table 2).

**Table 2.** Electrode coverage across patients. Included patients experimental IDs (N=14), and number of electrode contacts for each mesial temporal anatomical region, and 7 canonical networks using the Yeo-parcellation. The number of sites used for analysis is displayed. AMY: amygdala; HC: hippocampus; PHG: parahippocampal gyrus; VIS: visual (Yeo 1); SM: somatomotor (Yeo 2); DA: dorsal attention (Yeo 3); VA: ventral attention (Yeo 4); LIM: limbic (Yeo 5); FP: frontoparietal (Yeo 6); DM: default mode (Yeo 7).

| Patient | AMY | HC | PHG | VIS | SM | DA | VA | LIM | FP | DM |
| --- | --- | --- | --- | --- | --- | --- | --- | --- | --- | --- |
| S01 | 2 | 5 | 0 | 17 | 53 | 16 | 33 | 35 | 25 | 43 |
| S02 | 0 | 0 | 0 | 8 | 21 | 19 | 14 | 0 | 20 | 27 |
| S03 | 2 | 1 | 6 | 5 | 36 | 9 | 17 | 23 | 18 | 46 |
| S04 | 3 | 4 | 0 | 10 | 35 | 14 | 14 | 18 | 50 | 51 |
| S05 | 3 | 3 | 4 | 2 | 59 | 9 | 22 | 27 | 19 | 73 |
| S06 | 2 | 1 | 0 | 0 | 3 | 1 | 11 | 0 | 37 | 36 |
| S07 | 0 | 0 | 0 | 0 | 12 | 4 | 14 | 19 | 57 | 34 |
| S08 | 8 | 10 | 3 | 3 | 31 | 4 | 12 | 63 | 5 | 36 |
| S09 | 2 | 0 | 6 | 3 | 41 | 7 | 16 | 20 | 32 | 26 |
| S10 | 7 | 4 | 3 | 6 | 47 | 5 | 23 | 49 | 42 | 46 |
| S11 | 14 | 1 | 5 | 2 | 29 | 1 | 15 | 35 | 20 | 52 |
| S12 | 3 | 11 | 5 | 35 | 22 | 38 | 9 | 9 | 22 | 31 |
| S13 | 3 | 2 | 0 | 16 | 17 | 6 | 12 | 18 | 14 | 19 |
| S14 | 8 | 6 | 0 | 4 | 35 | 2 | 12 | 10 | 28 | 27 |

### Signal pre-processing

Recording sites in the reported SOZ (*n*=145) and those exhibiting epileptiform activity (*n* = 430, seizure or interictal) at any time during the paradigm recording were excluded. Channels were further rejected if they exhibited voltage values exceeding 5 standard deviations in more than 5% of the samples. Out of the 2,415 recorded electrode contacts across patients, a total of 2310 remained for further analyses, resulting in the mesial and Yeo7 network-parcellation electrode coverage (number of contacts) shown in Table 2. Remaining iEEG signals were denoised using the demodulated band transform (DBT) method^32^.To deal with high frequency noise common across channels we applied SVD (singular value decomposition) to the high-pass (250 Hz) filtered signals of all recorded channels and the first component was further extracted.

### Ripple detection

For the purpose of ripple detection, iEEG signals were bipolar-referenced to white matter^20,33^. Our rationale for using white matter activity as re-referencing procedure is that white matter presents lower intrinsic electrical activity as compared to the gray matter, while also carrying common-mode artifacts. Furthermore, similar to standard bipolar montages, white matter reference electrode contacts within the same grid or depth electrode often share similar impedance and noise profiles with their gray matter counterparts. This makes white matter electrode contacts suitable for reducing shared noise while preserving relevant signals (in our case, ripples).

White-matter (WM) signals were visually inspected to guarantee they did not exhibit pathological activity or residual noise. Mesial channels (HC, AMY and PHG) were re-referenced to the closest viable WM contact, preferentially in the same strip or electrode. Signals from other brain areas were re-referenced to an average signal calculated from non-mesial WM sites. Crucially, the WM channels used for HC re-referencing were not used to re-reference other signals. In order to verify the rejection of potential artifacts by WM iEEG referencing, we compared the detection of neural events after using common average and Gram-Schmidt referencing methods^34,35^ (Fig. S3A). Event detection by each referencing method was in good agreement (Fig. S3B, C), providing further confidence that the detected events reflect physiological activity, rather than noise arising from common sources.

After re-referencing, iEEG signals were filtered in the human ripple (70-160 Hz) frequency band ^20,36^ with a 4th order infinite-impulse response (IIR) Butterworth filter. This band-limited signal was then rectified, filtered below 25 Hz and z-scored. Candidate events were detected as epochs exceeding a threshold proportional to the standard deviation of the signal, calculated including the pre- and post-exercise periods (4 SD in this case). When the instantaneous power of the z-scored signal exceeded 20 SD, events were considered noise and were discarded. We compared the output of this standard ripple detection methodology to a non-negative matrix factorization (NMF) algorithm^18^ (Fig. S4) which offers a computationally efficient, and more unsupervised alternative. This method was applied to broadband spectral estimates (10-180 Hz) of the iEEG signals, yielding a low-rank decomposition of the peri-event spectra into a user-defined number of clusters. After applying this procedure, only events with a spectral support localized within the ripple band (70-160 Hz) were retained for cross-checking ripple detection. Both the standard and the NMF methodologies yielded similar event rates, with satisfactory mean event co-labeling proportion (83.91±5.47% mean with 95% confidence interval, Fig. S4). We additionally confirmed the reliability of our event detection in the hippocampus by visual inspection by experienced neurophysiologists.

### Ripple features

Three distinct features were computed for all detected ripple episodes (rate, duration and peak frequency). The rate estimate was computed as the total number of ripple events over the total recording time of each resting behavioral epoch (pre- or post-exercise). Ripple duration was computed as the full-width at half-maximum of the peri-event filtered (70-160 Hz) and rectified iEEG signal. Spectral analysis was performed using Morlet-wavelet spectrograms, unless otherwise specified. Spectrograms were z-scored with respect to baseline surrogate spectrograms computed using the same number of events with randomized inter-event intervals. Ripple peak frequency was then computed as the frequency corresponding to the maximum power of the Morlet-wavelet spectrogram, averaged in a time window of 50 ms centered at each event occurrence.

### Heart rate correlations

We first averaged the values of each ripple feature across contacts located in each region or network of interest. Then, we calculated the change in ripple features between the two resting state periods (Post Exercise – Pre Exercise). Thus, per subject, we had one single value that represents the average exercise-elicited change per region or network of interest during resting state. We then computed a Pearson correlation coefficient per region between such change in ripple features (rate, duration and peak-frequency) and each HR measurement (HR during pre-exercise RS, HR during the exercise session, HR during post-exercise RS, or HR difference between pre- and post-exercise RS). It is worth noting that in Fig. 3 we have one data point per subject. We excluded one subject from the correlation analyses (S04) as their HR measurements are outliers and we could not corroborate those measurements in the exercise and post-exercise period from the original source. For transparency, we depict the data point(s) corresponding to that subject in the corresponding plots with a cross.

### Statistical coupling between ripple time stamps

We estimated the statistical coupling between ripple events detected individually from HC and cortical iEEG signals. For that purpose, we defined hippocampal-cortical channel pairs using one ‘anchor’ hippocampal site per subject. When more than one HC site was viable, anatomical location was used as selection criteria. Specifically, HC sites contralateral to the SOZ and putatively closer to CA1/CA3 were preferred^17,20,37^.

Following ^38^ we obtained a histogram-type estimate of the cross-correlation functions of the (bivariate) point-processes corresponding to paired event types. For a set of ripple events occurring in the interval [0,*T*] of type *S*={*s_1_*,*s_2_*,…,*s_n_*} and type *K*={*t*_1_,*t*_2_,…,*t_m_*}, the cross-correlation estimates are based on counts in the following set:

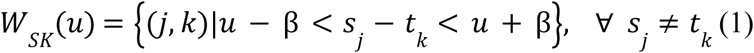

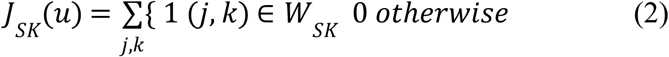

where 2β and *u* correspond to the histogram bin size and the center of the bin. Using *J_SK_*(.), we get the rate of coinciding events (in events/s) for a single experiment block as follows:

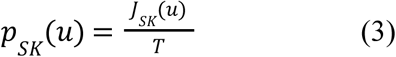

Statistical testing for the significance of the ripple coupling between areas was assessed by generating null-hypothesis distributions of the cross-correlation values in the time window of ±250ms around zero lag. These correlation values were computed from 400-point processes of the same rate, drawn at random from a uniform distribution. The ground-truth cross-correlation values were then compared with that of the null-hypothesis distributions, from which a *p*-value was computed ^37^. Correlation values of *p*<0.01 were considered statistically significant. For the group-level statistical analysis (LME model), to avoid effects induced by baseline correlation values, cross-correlograms were baseline-corrected using the average of the above null-hypothesis distribution across time lags.

### Amplitude-weighted phase locking (awPLV) analysis

Amplitude-weighted phase locking (awPLV) is an estimate based on the cross-spectrum *S_xy_* between two signals *x* and *y*. In order to factor out the influence of amplitude covariance, the cross spectrum is normalized with 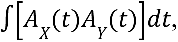 where *A_X_* (t) and *A_Y_* (t) correspond to the instantaneous amplitudes of the signals *x* and *y* in a time window around the event occurrence. This normalization results in a weighted-average phase coherence (hence the name “amplitude-weighted”):

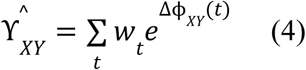

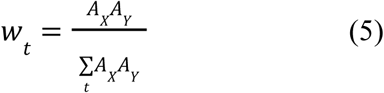

Like the standard PLV, the magnitude of this measure will be 1 if the signals *x* and *y* are perfectly phase locked, regardless of their amplitudes. In the absence of locking, the expected value of PLV is related to effective sample size ^39^. Thus, in order to compute pair-contact averages (here, across detected ripple events), effective sample size was approximated as:

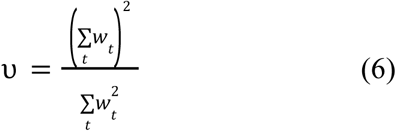

with sample bias:

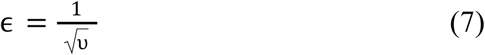

Therefore, the sample bias-corrected awPLV measure writes:

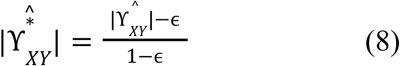

Spectral estimates of this analysis were performed using the demodulated band transform (DBT) across mesial-cortical contact pairs for frequency values below 200 Hz. As for the coupling (cross-correlation) analysis, we used one ‘anchor’ hippocampal contact per subject. Peri-ripple magnitude awPLV was computed in a time window of ±250 ms around the occurrence of hippocampal ripples to account for the trade-off between spectral and temporal resolution of the spectral estimate^32^. Estimates of awPLV across detected events were then averaged. Finally, an average of the resulting awPLV in the ripple frequency band (70-160 Hz) per channel pair was then used for further statistical analysis across brain regions and subjects (see Mixed-effects statistical model). Only for visualization purposes in Fig. 5B, sample bias-corrected awPLV was computed using complex Morlet-wavelet time-frequency spectrograms.

### Statistical Analysis

Effects of exercise on ripple features, ripple coupling, and peri-ripple awPLV were examined separately. To this end, we used linear mixed-effects models (LMEM). This method provides a principled way to account for both participant-level variability and heterogeneity in electrode coverage, while testing for region-specific effects of exercise on ripple dynamics. By modeling subject- and channel-level random effects, the approach increases robustness and generalizability of the findings despite the constraints of our intracranial datasets. Each model included experimental epoch (pre-, post-exercise) and anatomical region of interest (ROI) entered as within-subject factors. Random effects included an intercept and epoch effect (slope) modeled by subject ID (1 + epoch | SubjectID) and channel ID nested within subject ID (1 + epoch | SubjectID:ChanID), which accounted for variability unique to participants and their specific recording montage. Fixed effects of interest included the interaction between epoch and ROI for each level of the ROI factor (e.g., epoch:ROI interaction). Statistical significance was determined based on an interaction term having a *p*-value of *p*<0.05 within each model. Multiple statistical tests (e.g. across networks) were corrected for multiple comparisons when indicated. In the mesial temporal parcellation, we defined three anatomical ROIs: HC, AMY, and PHG. For the neocortical parcellation we used seven canonical networks^33^: Visual (VIS), Somatomotor (SM), Dorsal Attention (DA), Ventral Attention (VA), Limbic (LIM), Frontoparietal (FP), Default-mode (DM) network (Fig 1B).

## Results

### Physical activity and exercise

Participants (*N*=14, 10 males) were patients with drug resistant epilepsy being monitored for epileptic seizure localization with chronic intracranial electroencephalography (iEEG) recordings. Patients completed an acute exercise paradigm while neural activity was continuously monitored. The paradigm consisted of a pre-exercise resting state period (∼20 min), followed by a period of guided exercise on a pedal exerciser (∼20 min), and then by a post-exercise resting state period (∼20 min) (Fig. 1A). During resting periods, participants were instructed to relax with their eyes closed. During exercise, participants started with a warm-up (∼5 min), and then cycled at a target range of light-to-moderate intensity aerobic exercise [50-60% of their age-predicted maximum heart rate (HR%)]. Participant demographics and physiological response to exercise are summarized in Table 1. Participants (median age = 23.5, range 17 to 50) reported an average of 7.6 hours (*SD* = 3, min-max 3-14) of non-sedentary physical activity per day, including both light and moderate-to-vigorous intensity activity. Compliance to the exercise protocol was satisfactory, with an average HR% during exercise of 57.7% (*SD* = 5.7), with most participants within or near the target range and as a group were elevated compared to the pre-exercise rest (*t*(12)=7.12, *p*<0.001; Fig. 1A). Similarly, self-reported RPE was an average of 11.9 (*SD*=1.5), corresponding to a subjective intensity of “light” to “somewhat hard” on the scale matching their target intensity zone, and was significantly elevated compared to the pre-exercise rest (*t*(11)=7.57, *p*<0.001). Age, sex, and self-reported physical activity were not correlated with acute exercise intensity (*p*’s all >.05), and therefore were not entered as covariates in analyses reported in this study.

### Electrode coverage and neural signals

Electrode coverage across patients is summarized in Fig. 1B. Signals from 2415 recording sites were collected from the 14 patients included in this study. Cortical recording sites were grouped into 7 canonical networks using the Yeo-parcellation^40^ according to their anatomical location. Contacts out of the brain, in the seizure onset zone (SOZ), or exhibiting interictal epileptiform discharges (IEDs; Fig. S1) during the recordings were excluded from the analysis. Automated noise rejection and denoising steps were subsequently applied to the remaining signals (Materials and Methods; Fig. S2). The findings described in the following sections result from the comparison between the pre- and post-exercise resting state periods, while subjects stayed quiescent, silent, and with closed eyes.

### Acute exercise modulates ripple properties

Ripples were reliably identified in the iEEG signals recorded from depth electrodes targeting HC^19,20^, PHG^36,41^, and AMY^42^ (Fig. 1C-E). Briefly, these oscillatory events were detected as peaks exceeding 4 SD threshold in the z-scored envelope of the iEEG signal filtered in the 70-160 Hz frequency range (Materials and Methods).

The rate of hippocampal ripples and their electrophysiological features are known to vary across behavioral states and to predict memory performance^43,44^. We hypothesized that exercise would be associated with an increase in hippocampal ripple occurrence and modulation of electrophysiological features. Fig. 2A shows exemplary raw traces of peri-ripple iEEG of HC, where variability in duration and power spectral density are apparent. To investigate whether exercise modulates ripple features, a LMEM was deployed separately for ripple rate, duration and peak frequency (Materials and Methods). We found statistically significant pre-to-post exercise ripple rate increases in the HC (*b*=0.06, *SE*=0.02, *t*(170)=3.02, *p*=0.002; Fig. 2B, left boxplot). However, exercise did not significantly affect hippocampal ripple duration or peak frequency (Fig. 2B, middle and right boxplots).

**Figure 2.**
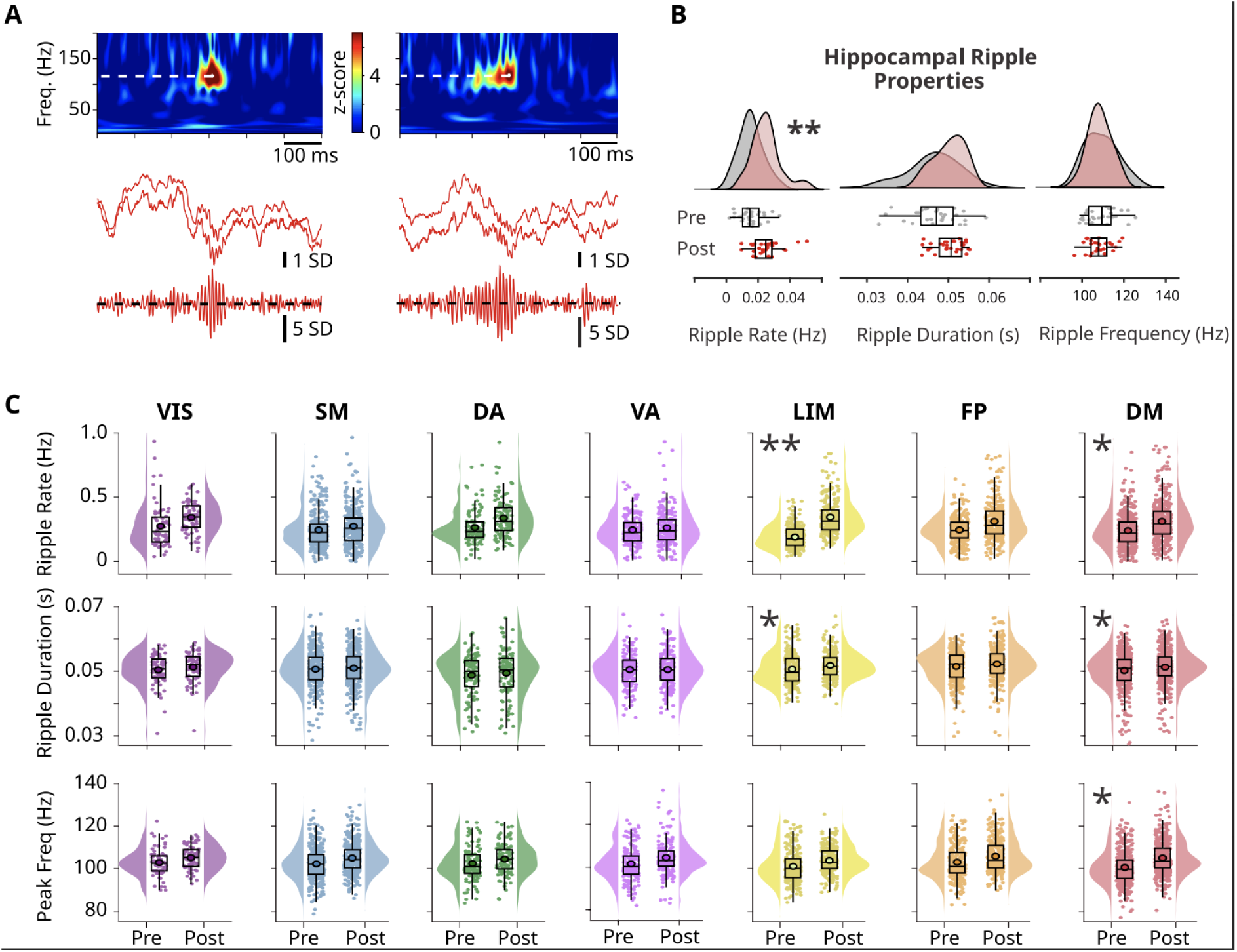
Effect of acute exercise on ripple properties. **(A)** Exemplary raw (<300 Hz) and filtered (70-160 Hz) peri-ripple iEEG traces with spectrograms showing that ripples in the human hippocampus vary in duration and peak frequency. White, dotted lines in the spectrograms indicate ripple frequency peak at time zero. Two traces of the raw iEEG per anatomical structure are displayed to show the variability across contiguous sites. **(B)** Across-subjects distributions of ripple rate, duration and peak frequency in the resting state pre- and post-exercise in the hippocampus (HC), and **(C)** in the 7 canonical networks, illustrating the modulation of ripple characteristics by acute exercise. In box plots, lines crossing the boxes indicate the median, box edges indicate 25th and 75th percentiles, and data points beyond the whiskers are outliers. Each dot represents one recording site. \**p*<0.05, \*\**p*<0.01 according to a LMEM treating subject and recording sites as random effects.

In light of these observations, we investigated whether the electrophysiological features of ripple events were also modulated in the rest of the brain as a putative consequence of acute physical activity. To examine this possibility, after ripple detection and feature computation, cortical sites were grouped according to seven canonical networks^31^. Between-subjects statistical analysis was again performed using the same LMEM, which treated subjects and recording sites as random effects. Consistent with our previous results, exercise increased ripple event rate and duration in the limbic network (rate: *b*=0.065, *SE*=0.014, *t*(3330)=4.59, *p*=0.00000441; duration: *b*=0.000545, *SE*=0.000256, *t*(3327)=2.12, *p*=0.0000426; Fig. 2C). Notably, the DMN exhibited significant exercise-related increases in ripple rate, duration, and peak frequency (rate: *b*=0.030, *SE*=0.013, *t*(3330)=2.23, *p*=0.025; duration: *b*=0.00047, *SE*=0.00023, *t*(3327)=1.99, *p*=0.045; peak frequency: *b*=1.84, *SE*=0.87, *t*(3327)=2.11, *p*=0.034). No other canonical networks showed exercise-related change in ripple features (Fig. 2C).

To further guarantee that these results do not rely on pathological activity, we repeated the LMEM analyses using only the contacts from subjects with bilateral coverage (*N* = 6) from the hemisphere contralateral to the SOZ. In this subset of contacts our main results remained consistent, reaching higher statistical significance in some cases (Fig. S5).

These results indicate that acute exercise elicits an increase in the rate of ripple events and modulates their properties (duration and peak frequency) in limbic areas and the DMN.

### Modulation of resting-state ripple properties correlates with heart rate during exercise

We investigated whether and how HR during exercise predicts the observed changes in resting state ripple rate and ripple electrical properties. To this end, we performed correlation analyses (Materials and Methods). We found positive correlations between the HR during exercise and the subsequent change in resting state ripple rate (i.e., from the pre-exercise to the post-exercise periods). This was the case for the DM (*r*=0.68, *p*=0.01), the VA (*r*=0.67, *p*=0.01) and the FP (*r*=0.65, *p*=0.02) networks (Fig. 3A). In contrast, neither the HR values in the pre- or post-exercise period, nor the difference between them, were significantly correlated with ripple rate or properties in any brain network. Next, we explore whether exercise elicited changes in hippocampal-neocortical interactions.

**Figure 3.**
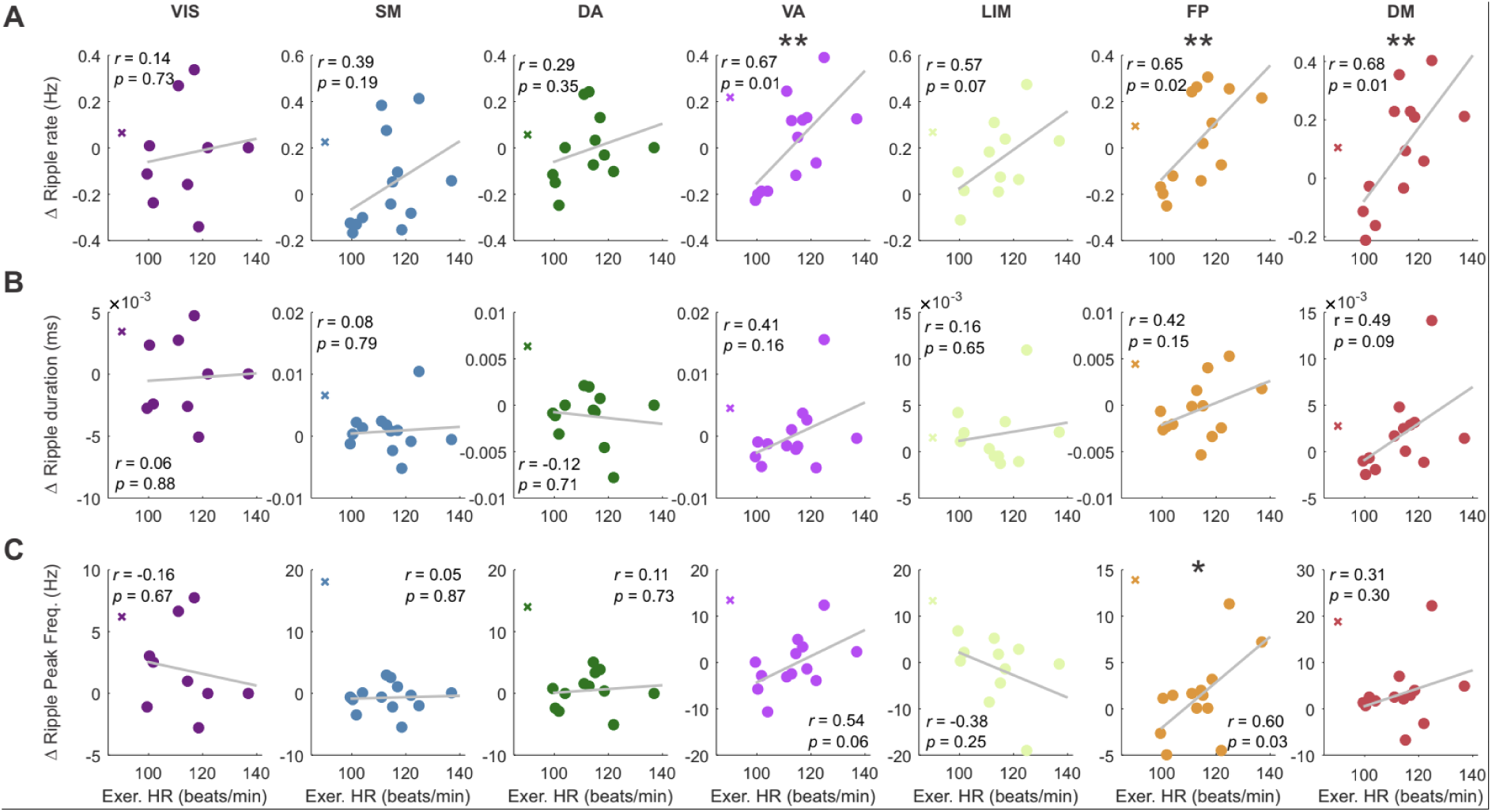
Modulation of neocortical ripple properties is correlated with exercise-associated heart rate across subjects. Across-subjects scatter plots in the 7 canonical networks illustrating the correlation between change in ripple properties (RS PostExercise – RS PreExercise) and HR during the exercise period. **(A)** Ripple rate, **(B)** Ripple duration, and **(C)** Ripple peak frequency. Each dot corresponds to one subject. The subject marked by “X” was depicted in the plot for transparency but excluded from the calculation as it represents an outlier and we were not able to verify the measurements from the source records. Asterisks indicate statistically significant correlations, \**p*<0.05, ** *p*<0.001. In contrast, neither the HR values in the pre- or post-exercise resting state periods, nor the difference between them were significantly correlated with ripple rate or electrical properties.

### Acute exercise increases hippocampal-cortical ripple coupling

Ripples in mesial areas often co-occurred with similar short-lived high-frequency events in other brain areas during pre- and post-exercise resting state (Fig. 4 and Fig. 5; see also Fig. S1). To test whether exercise modulates ripple hippocampal-cortical coupling, we performed cross-correlation analyses between the time stamps of ripples in the hippocampus (anchor events) and other sub-cortical and cortical sites. To this end, we paired one hippocampal recording site per subject (selected based on its anatomical location) with the other sites of interest. To guarantee an unbiased coupling estimate, cross-correlation values per pair were corrected for potential ripple-rate baseline effects by subtracting baseline values obtained by a permutation procedure (Materials and Methods).

**Figure 4.**
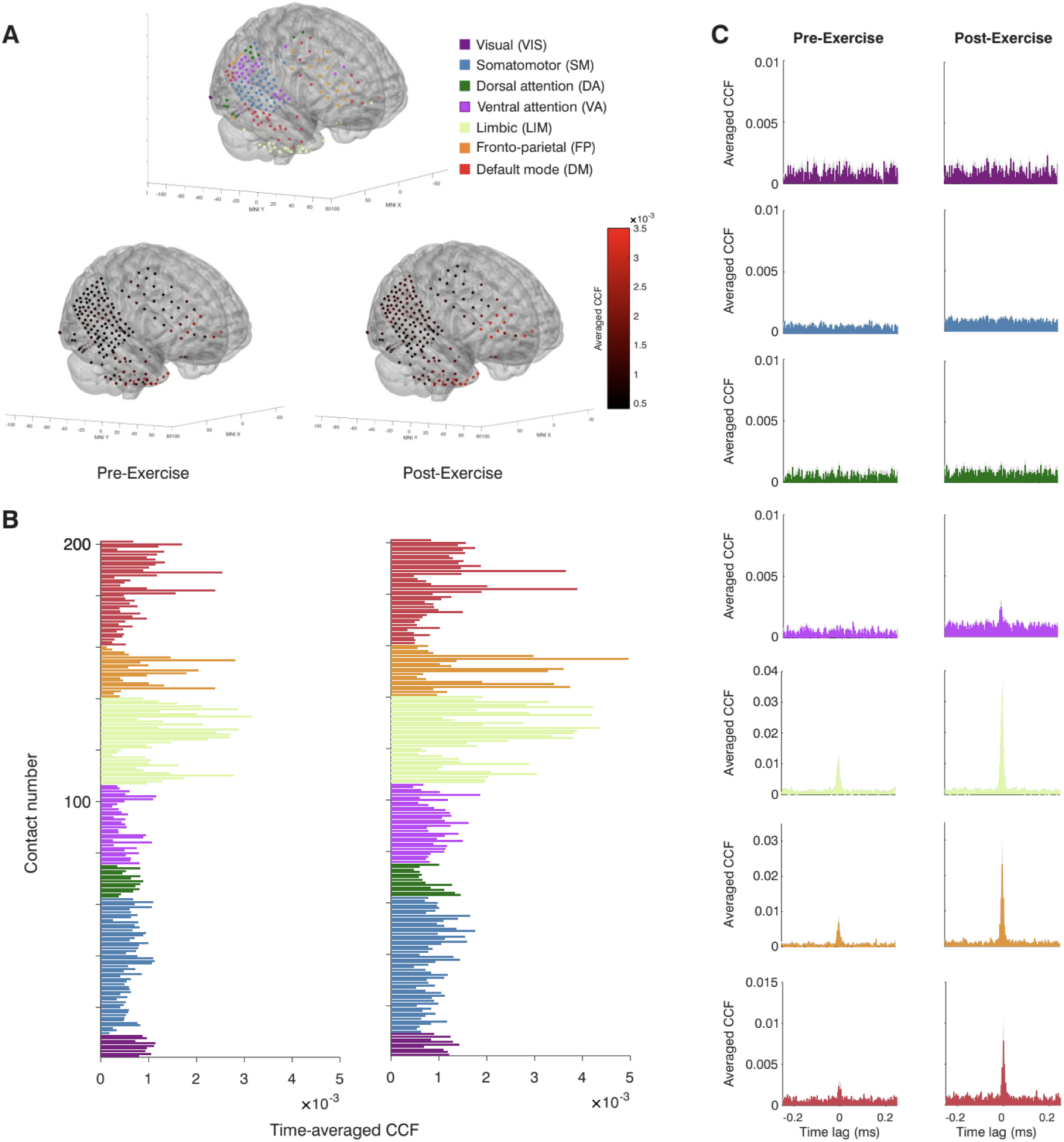
Hippocampal-cortical ripple statistical coupling in an exemplary subject. **(A)** Recording contacts’ distribution in the neocortex classified into 7 canonical networks according to their anatomical locations (top). Pre- (bottom, left) and post-exercise (bottom, right) resting state cross-correlation values between hippocampal ripples and ripples detected across all cortical contacts, overlaid in the same brain template. **(B)** Same as (A), but the averaged pre- (left) and post-exercise (right) resting state cross-correlation values are presented as bars, and sorted according to the 7 canonical networks. In this subject there is overall larger coupling between hippocampus and the VA, limbic, FP and DM networks during post-exercise resting state. Note that across subjects only limbic, DM and VA reached significance. **(C)** Cross-correlograms between hippocampal and cortical ripples, averaged across cortical contacts associated with each canonical network. Exercise-induced hippocampal-cortical ripple coupling effects were observed in networks 4 to 7 (VA *p*=4.6e-07, LM *p*=7.66e-07, FP *p*=4.00e-05 and DM *p*=7.84e-07, Bonferroni-corrected sign test). Shadings indicate standard error of the mean (SEM).

**Figure 5.**
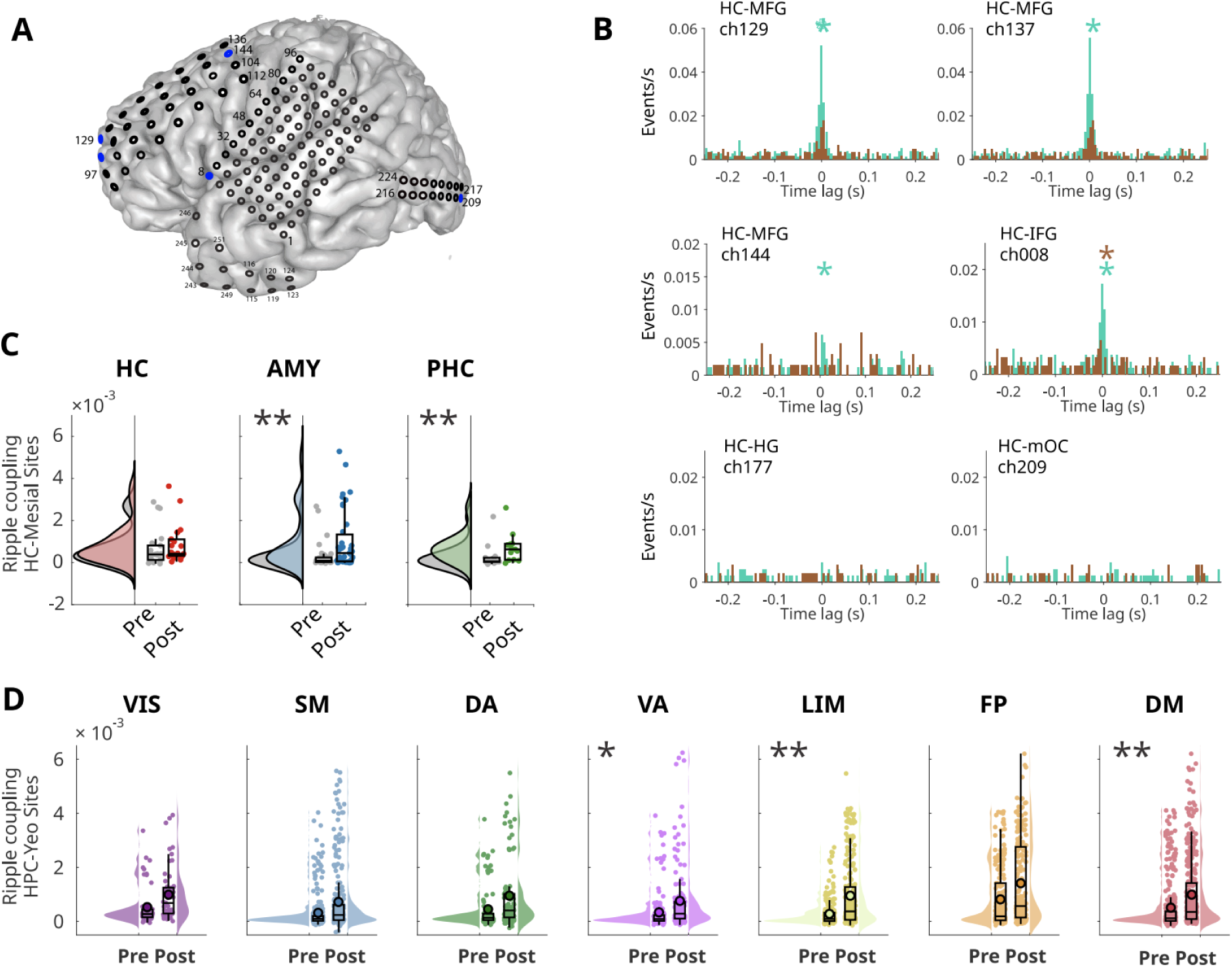
Effect of exercise on hippocampal-cortical peri-ripple coupling. **(A)** Recording contacts’ distribution in the neocortex of the subject displayed in Fig. 4. Blue dots indicate contacts shown in panel B. **(B)** Exemplary cross-correlograms between hippocampal ripples and ripples from distinct extra-hippocampal sites (MFG: middle frontal gyrus; IFG: inferior frontal gyrus; HG: Heschl’s gyrus; mOC: middle occipital gyrus). Asterisks indicate significant coupling around lag zero (*p*<0.01, permutation t-test). **(D)** Across-subjects pre- and post-exercise resting state coupling values between hippocampal ripples and ripples detected in other mesial areas (i.e., HC, AMY, and PHG). **(E)** Across-subjects pre- and post-exercise resting state coupling values between hippocampal ripples and ripples detected across 7 canonical networks: visual (VIS), somatomotor (SM), dorsal attention (DA), ventral attention (VA), limbic (LIM), frontoparietal (FP) and default-mode (DM). **(D** and **E)** LME models for the comparison between pre- and post-exercise resting state blocks. In box plots, lines crossing the boxes indicate the median, box edges indicate 25th and 75th percentiles, and data points beyond the whiskers are outliers. Asterisks indicate an statistically significant effect, \**p*<0.05, ** *p*<0.001.

Hippocampal-cortical ripple statistical coupling is significant, but occurs selectively across cortical channels (Fig. 4A). Fig. 4B shows pre- and post-peri-event zero-lag correlation values (Materials and Methods) across all hippocampal-cortical channel pairs in an exemplary patient. The average hippocampal-cortical ripple cross-correlation across the seven canonical networks in this exemplary patient are shown in Fig. 4C, with exercise-induced hippocampal-cortical ripple coupling effects occurring in VA, LIM, FP and DM (VA *p*=4.6e-07, LIM *p*=7.66e-07, FP *p*=4.00e-05 and DM *p*=7.84e-07, Bonferroni-corrected sign test).

As in the previous section, the influence of exercise in hippocampal-other mesial areas ripple coupling, and hippocampal-cortical ripple coupling at the population level was tested using a LMEM. In this case, the dependent variable was the average cross-correlation around ripple occurrence in the hippocampus (Materials and Methods; Fig. 5A-B). We found that exercise significantly increased HC-AMY (*b*=0.000342, *SE*=0.0000878, *t*(144)=3.89, *p*=0.00015; Fig. 5C, middle subpanel), and HC-PHG ripple coupling (*b*=0.000296, *SE*=0.000106, *t*(144)=2.78, *p*=0.006; Fig. 5C, right subpanel).

Hippocampal-cortical group-level LMEM-based analysis shows that, after exercise, ripple coupling increases between the HC and the limbic network, and between the HC and DMN (limbic: *b*=0.000242, *SE*=0.000103, *t*(2752)=2.34, *p*=0.019; DMN: *b*= 0.000231, *SE*=0.000102, *t*(2752)=2.26, *p*=0.023; Fig. 5D). Additionally, the VA network exhibits enhancement in coupling with hippocampal ripples (*b*=0.000219, *SE*=0.000104, *t*(2752)=2.09, *p*=0.035). In contrast, the effect was not significant in the remaining four networks (VIS, SM, DA and FP, *p*>0.05). These results were consistent across peri-ripple windows of different lengths, confirming the robustness of the effect in the limbic and DM networks. Thus, hippocampal ripples selectively co-occur with ripples in these networks, and this coupling is larger during post-exercise resting state, as compared to pre-exercise resting state. Altogether, our results indicate that exercise not only modulates hippocampal ripple properties, but also enhances hippocampo-neocortical and intra-mesial ripple coupling.

### Hippocampal-cortical ripple phase synchrony is enhanced by exercise

Recent studies have shown that high frequency oscillations are synchronized between distributed brain regions and reliably exhibit a network organization^21,22^. As ripples are a manifestation of volleys of spiking neurons, ripples (and hence population spike discharges) may be phase-locked across distinct, yet functionally connected brain areas. Given the increase in ripple coupling we observed after exercise, we explored whether exercise also modulated ripple phase synchronization.

To this end, the time stamps of ripples detected in the hippocampus were used as triggers to build peri-ripple iEEG traces across all recorded sites. The phase synchrony between hippocampal and cortical ripples was measured by computing an amplitude-weighted phase-locking value (awPLV). Briefly, this quantity measures the degree to which two oscillatory signals are phase-locked, regardless of their amplitude^39^ (Materials and Methods). Ripple-triggered awPLV responses were computed separately for pre-exercise and post-exercise resting state blocks in a ±250 ms window around the events’ occurrence, the averaged awPLV in the ripple frequency band (70-160 Hz) was then used for statistical analysis (Fig. 6A-B displays this quantity for an exemplary subject; see also Fig. 7A). After exercise, hippocampal-cortical ripple phase coupling increases selectively in the ripple frequency band for one exemplary patient in Fig. 6C (phase locking peaks around 100 Hz). This modulation is also network-specific; for example, in the exemplary patient, hippocampal-cortical ripple coupling effects occurred significantly in LIM, FP and DM networks (LIM *p*=2.85e-08, FP *p*=0.01, and DM *p*=0.003, Bonferroni-corrected sign test).

**Figure 6.**
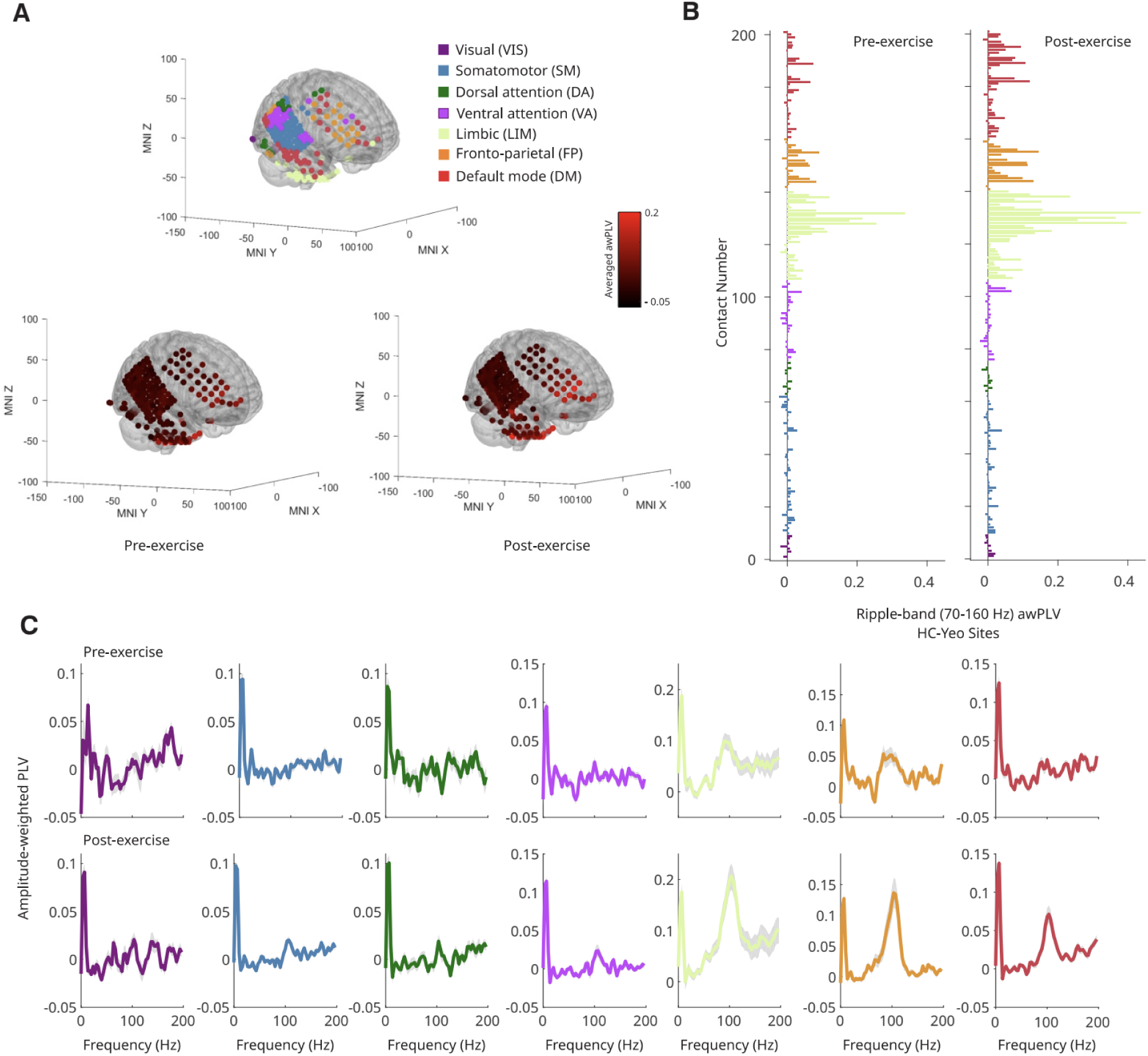
Hippocampal-cortical ripple phase synchrony in an exemplary subject. **(A)** Recording contacts’ distribution in the neocortex classified into 7 canonical networks according to their anatomical locations (top). Pre- (bottom, left) and post-exercise (bottom, right) resting state amplitude-weighted phase locking value (awPLV) in the ripple frequency band (70-160 Hz) between hippocampal and neocortical peri-event activity, overlaid in the same brain template. **(B)** Same as (A), but the averaged pre- (left) and post-exercise (right) resting state awPLV in the ripple frequency band is presented as bars, and sorted according to the 7 canonical networks. In this subject there is overall larger ripple phase coupling between hippocampus and the LIM, FP and DM networks during post-exercise resting state. **(C)** awPLV between hippocampal and cortical ripples estimated via demodulated band transform (DBT). Values were averaged across cortical contacts associated with each canonical network. Exercise-induced hippocampal-cortical ripple coupling effects were observed in networks 5 to 7 (LM – *p*=2.85e-08, FP – *p*=0.01, and DM – *p*=0.003, Bonferroni-corrected sign test). Shadings indicate standard error of the mean (SEM).

**Figure 7.**
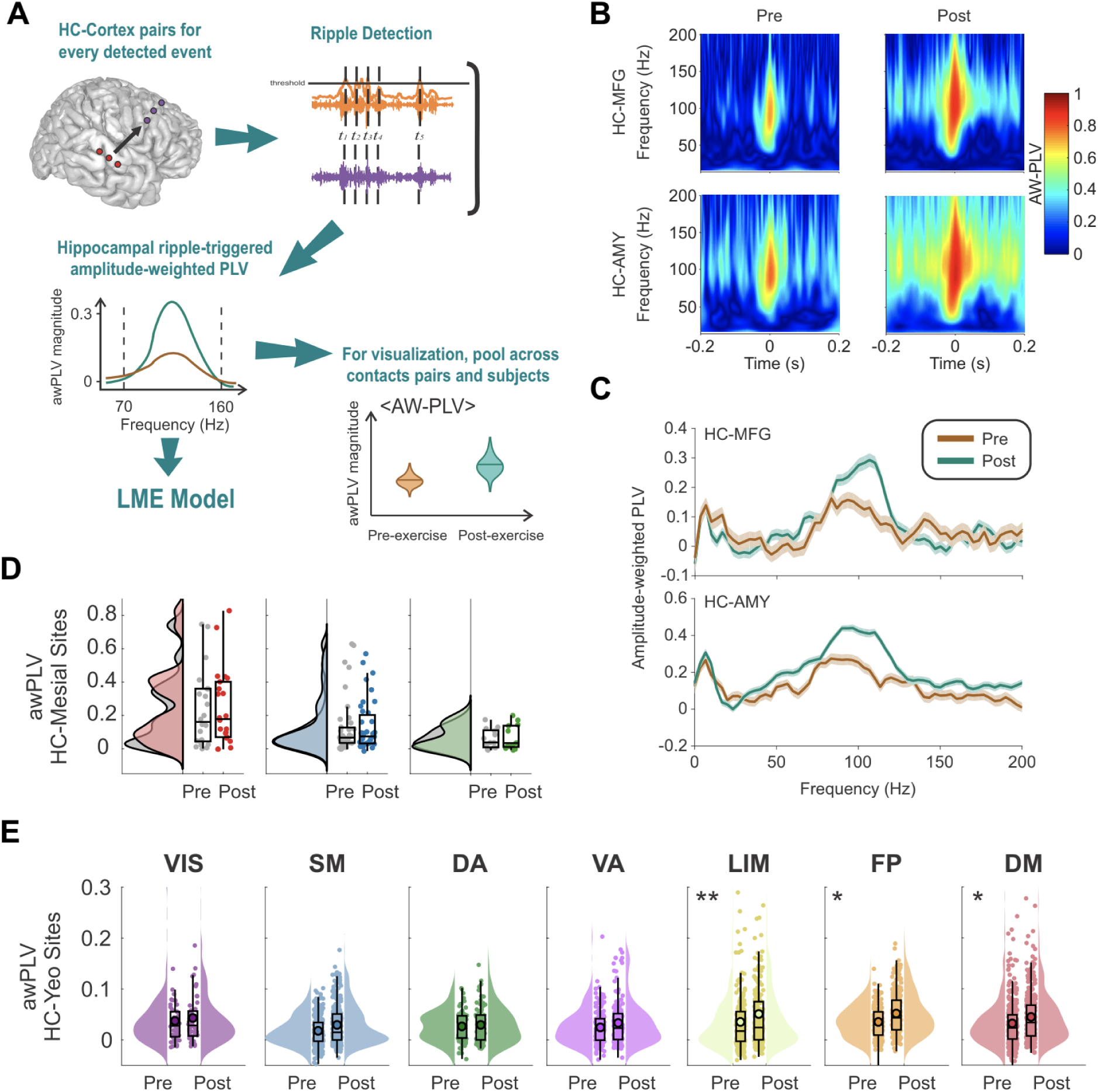
Effect of exercise on hippocampal-cortical peri-ripple phase synchrony. **(A)** For phase synchrony calculation, hippocampal ripple timestamps are used to compute a peri-event amplitude-weighted phase locking value (awPLV) between hippocampal-cortical site pairs. The ripple band (70-160 Hz) averaged awPLV estimate per site pair is then used for group-level statistical analysis using a LMEM. **(B)** 2-D awPLV estimated from complex Morlet-wavelet spectrograms between the hippocampus and middle frontal gyrus (MFG) (top), and between the hippocampus and amygdala (bottom), shows that awPLV between hippocampus and other brain areas increases during post-exercise (right), relative to pre-exercise resting state (left). **(C)** Same as panel (B), but the awPLV estimate is 1-D and calculated via demodulated band transform (DBT). **(D)** Across-subjects pre- and post-exercise resting state awPLV at the time of hippocampal ripples between hippocampus and other mesial areas (i.e., HC, AMY, and PHG). **(E)** Across-subjects pre- and post-exercise resting state awPLV at the time of hippocampal ripples, between hippocampus and 7 canonical networks: visual, somatomotor, dorsal attention, ventral attention, limbic, frontoparietal and default-mode. **(D** and **E)** LME models for the comparison between pre- and post-exercise resting state blocks. In box plots, lines crossing the boxes indicate the median, box edges indicate 25th and 75th percentiles, and data points beyond the whiskers are outliers. \**p*<0.05, \*\**p*<0.001.

To illustrate that the awPLV increases are localized in frequency and peri-ripple time, the averaged HC-MFG time-frequency awPLV for an exemplary contact pair (one MFG contact belonging to the DMN) is shown in Fig. 7B (in line with the 1D estimates of Fig. 6, and Fig. 7C). Notably, consistent with Fig. 6, post-exercise peri-ripple awPLV was larger than its pre-exercise counterpart, indicating a higher ripple-band phase coupling of the former with respect to the latter. Analogously to the former ripple cross-correlation analysis, statistical comparison at the group level between pre- and post-exercise blocks over cortical areas was performed using a LMEM to account for differences in participant electrode coverage. The dependent variable of this analysis was the pre- and post-exercise awPLV (Materials and Methods).

Whereas no significant effects were found for other mesial areas (Fig. 7D), we found that the limbic, FP, and DM networks showed a significant effect for awPLV across subjects (Fig. 7E; LIM: *b*=0.0085, *SE*=0.0027, *t*(2754)=3.11, *p*=0.001; FPN: *b*=0.0062, *SE*=0.0027, *t*(2754)=2.24, *p*=0.024; DMN: *b*=0.0056, *SE*=0.0027, *t*(2754)=2.09, *p*=0.036). These results indicate that hippocampal and neocortical high-frequency ripples are phase locked, and this coupling is modulated by exercise in some canonical networks but not others, with the largest effect in the limbic network.

## Discussion

In this investigation, we examined how acute exercise modulates hippocampal ripple properties and hippocampal-cortical ripple dynamics. We found that, after a session of acute exercise, ripples occur at higher rates in the hippocampus, and limbic and DM networks. We also found positive correlations between HR during exercise and exercise-induced change in resting state cortical ripple rate in several cortical networks including the DM, possibly pointing to a link between ripples^21^ and sympathetic response to physical exercise. Furthermore, we quantified the coupling between ripples in the hippocampus and ripples in neocortical sites, before and after the exercise session. Whereas hippocampal and neocortical ripples were both temporally- and phase-coupled in the pre-exercise resting state period, physical exercise enhanced the extent of this coupling, predominantly in the limbic and DM networks.

### Post-exercise modulation of ripple features

The rate of hippocampal ripples and their physiological features are known to vary across states and correlate with memory performance^43,44^. For example, in rodents, ripple duration increases as a result of mnemonically demanding situations, such as spatial learning^43^. Moreover, animal studies have also shown that aging is associated with reduced ripple rate during sleep and awake resting state, which may contribute to age-associated memory impairment^45^. In the context of these studies, an increase in hippocampal ripple rate elicited by exercise could explain, at least partly, the beneficial effects of physical activity in memory function. Our results advance the hypothesis of hippocampal ripples as a mechanism through which physical activity bouts may counteract age-related memory impairments. Future investigations could address whether and how these exercise-induced acute modulations contribute to long-term changes in neural plasticity.

### Exercise effects on hippocampal-cortical interactions

In line with animal studies^21,46^, we showed that hippocampal ripples co-occur with ripples in neocortical and other mesial areas. Mesial areas are functionally and anatomically connected with other structures of the limbic network (e.g. inferotemporal, orbital gyri)^47–49^ and associative cortices^16,21,31,50^. For example, hippocampal activity propagates to distinct neocortical targets following excitation of the output layers of the entorhinal cortex (EC)^51^, and/or via retrosplenial cortex^46^. Furthermore, EC and neocortical activity may also modulate mesial activity across several brain states^49,52–55^.

Furthermore, our results contribute to the growing literature showing that ripple coupling is a widely distributed phenomenon. Ripples occur simultaneously and phase-lock across multiple brain regions, even between hemispheres^22,56^. In the present study, across-subject comparisons between pre- and post-exercise epochs pointed to the limbic and the DM networks as consistently impacted by physical exercise. In this respect, we suggest that ripple dynamics may underlie the changes in hippocampal, limbic, and DMN functional connectivity observed in fMRI studies^57^.

The DMN is classically considered a resting-state network that supports inwardly oriented attention during episodic memory and future planning, as opposed to sensory-driven processing. Furthermore, non-human primate studies have shown that endogenous resting state DMN fluctuations detected in BOLD-fMRI correlate with the occurrence of hippocampal ripples^16,17,58^. In humans, the strongest ripple-coupled activations during autobiographic mnemonic recollection were found in the DMN^20^. In line with this evidence, it has been hypothesized that the DM^59^ and FP^60^ networks could trigger high-frequency oscillation (HFO) cascades spreading the reactivation of mnemonic and other representational content (e.g. semantic content)^59^. Altogether, this evidence advances a view of the DMN as a central hub in a bursting network (high synchrony neural events) that spreads information, possibly as reactivation or replay, across broad cortical areas^20,22,59,60^.

While enhanced post-exercise ripple coupling in specific brain networks suggests a potential positive effect of exercise on mnemonic processing, this hypothesis demands further investigation. Our study provides a first description of how physical activity influences hippocampal-cortical ripple dynamics, but does not directly test the deployment of exercise-induced ripples in cognition. Future studies should explore the connection between exercise-induced changes in hippocampal ripples and subjects’ performance in specific mnemonic tasks known to improve after acute exercise^61^.

### A neural substrate linking body state and cognition

Besides their role in cognition, hippocampal ripples play a role in glucose regulation and interoceptive signaling between the brain and the body. Specifically, in freely behaving rats, bursts of clustered hippocampal ripples are followed by decreased peripheral interstitial glucose after about 10 minutes, with ripple rate having the most impact on glucose compared to ripple frequency and duration^62^. Our finding that resting ripple rate increases in the human hippocampus after exercise presents an opportunity to test whether and how human hippocampal-cortical network ripple dynamics relates to reductions in peripheral glucose within and across individuals following exercise. Experimental conditions such as a fasted state and real-time measurement of peripheral glucose are feasible in this patient population^63^. Whereas the results of the present study suggest that the modulation of ripple dynamics is a mechanism by which exercise affects cognitive function, the role of metabolic changes (i.e. glucose) deserves further investigation and discussion. Crucially, the dual cognitive and metabolic roles of ripples are complementary rather than mutually exclusive. Ripples are then proposed as a metabolic marker embedded in hippocampal function, enabling the use of past experiences –stored in memory– to anticipate future metabolic demands according to predicted stimuli and planned behavior^64,65^.

Finally, further experiments are required to disentangle how exercise-associated metabolic changes account for the observed ripple effects, and to investigate how those bodily factors contribute to cognitive performance.

### Limitations of this study

Whereas our study provides novel mechanistic insights into how acute exercise modulates hippocampal-cortical communication in humans, it faces challenges common to invasive neurophysiological studies (iEEG) in human patients^19,33,36,66,67^. Given the size of the participant cohort (*N*=14) and the inclusion of patients with drug resistant epilepsy, we caution against overinterpretation of our findings. In this type of study, electrode coverage is determined by individual clinical needs and therefore varies across subjects. Whereas our analysis pipeline accounts for many of these factors (e.g. exclusion of SOZ contacts, signal denoising, LMEMs etc.), these methodological constraints remain important to acknowledge. Furthermore, although we report the results of our statistical models across canonical networks with uncorrected *p*-values, it is worth emphasizing that the observed neural activity patterns and exercise effects were consistent across patients, brain regions and measures, and thus unlikely to reflect chance findings.

Finally, the spatial resolution of extracellular field potentials depends on electrode type and electrical properties (e.g., impedance). Most intracranial clinical electrodes still yield relatively coarse field measurements, yet can capture high-frequency ripple activity. Importantly, electrode diameter influences impedance and thus the ability to detect high frequencies^68^, which constrains the resolution achievable in human iEEG compared to microelectrode recording methodologies. In addition, the specifics of human anatomy render the access to deep mesial structures such as the amygdala and the hippocampus cumbersome. Hippocampal depth electrode contacts must target the *cornu ammonis* (or nearby hippocampal substructures) in order to reliably detect *bona fide* sharp wave-ripple activity. The curved anatomical organization of the human *cornu ammonis* poses additional challenges for interpreting extracellular measurements and their potential translation to other experimental models, as such measurements in humans most likely reflect the summed contribution of cellular activities arising from both open- and close-field configurations^17,69^. In contrast, experimental animal models –specifically rodents– allow for precise and consistent targeting of hippocampal subregions in an open-field configuration. These models offer complementary opportunities to test the hypotheses investigated in this work, and those raised by our findings.

## Data availability

The data and code supporting the findings of this article will be available upon request to the lead contact, Prof. Dr. Michelle Voss.

## Acknowledgements

We acknowledge the generosity of the patients, who contributed time and effort to take part in this study.

## Funding

No funding was received towards this work.

## Competing interests

The authors report no competing interests.

## Notes

### Competing Interest Statement

The authors have declared no competing interest.

### Summary of Updates

The manuscript has been revised according to reviewers suggestions; for example, we added detail to the methods description and clarify our hypothesis. Scientific references have also been updated.

